# De-*N*-glycosylation of *in vivo* and *in vitro* adipogenic stem cell products unmasks differential expression of CD36 glycoprotein in human adipogenesis

**DOI:** 10.64898/2026.05.01.722121

**Authors:** Katherine Wongtrakul-Kish, Paul A. Haynes, Benjamin R. Herbert, Nicolle H. Packer

## Abstract

Adipogenesis is the process of adipose-derived stem cells (ADSCs) responding to extracellular signals from the stem cell niche to differentiate into adipocytes (fat cells) and may be studied *in vitro* using a cocktail of chemicals that promote adipogenic differentiation to produce differentiated ADSCs (dADSCs). The global membrane *N*- and *O*-glycosylation changes of this process have been previously analysed and compared to native adipocytes as a benchmark for a ‘true’ adipocyte profile, and revealed that bisecting GlcNAc type *N*-glycans are characteristic of adipogenesis. As stem cell differentiation has been widely reported to result in cellular protein changes, the same cells (ADSCs, dADSCs and mature adipocytes) were characterised for their membrane proteome here using label-free quantitative shotgun proteomics analysis. The membrane proteome displayed more differences in protein numbers between the cell types compared to the previously reported *N*-glycome which had shown high identical glycomes between stem cells and *in vitro* dADSCs, suggesting that the proteome is more dynamic during *in vitro* adipogenesis. Following the global shotgun proteomics analysis, a more targeted approach of carrying out proteomic analysis of de-*N*-glycosylated peptides of gel-separated proteins unearthed new glycoproteins not detected in the shotgun proteomic analysis. This approach identified the adipogenic marker, CD36, to be under-represented in the shotgun proteome analysis, but as the dominant (glyco)protein in the adipocyte membrane proteome that was also up-regulated at the mRNA transcript level in both the *in vitro* differentiated ADSCs (7.1-fold increase) and mature adipocytes (102.9-fold increase). A comparison of CD36 sequence coverage in the global shotgun analysis with the de-*N*-glycosylated CD36 revealed a 41% increase when *N*-glycans were removed prior to trypsin digestion, explaining its observed increased abundance and highlights the crucial need for de-*N*-glycosylation of proteins in proteomics experiments for increased identification of glycoproteins. The systems glycobiology approach by the integration of previously reported glycomics data and the proteomics and transcriptomics analyses in this work extended the investigation of membrane protein glycosylation changes in adipose-derived stem cell differentiation. The work provides a framework for future glycoproteomics-based investigations into the differentiation of stem cells into adipocytes, and will allow their related pathologies and potential therapeutic applications to be discovered.

**GRAPHICAL ABSTRACT:** 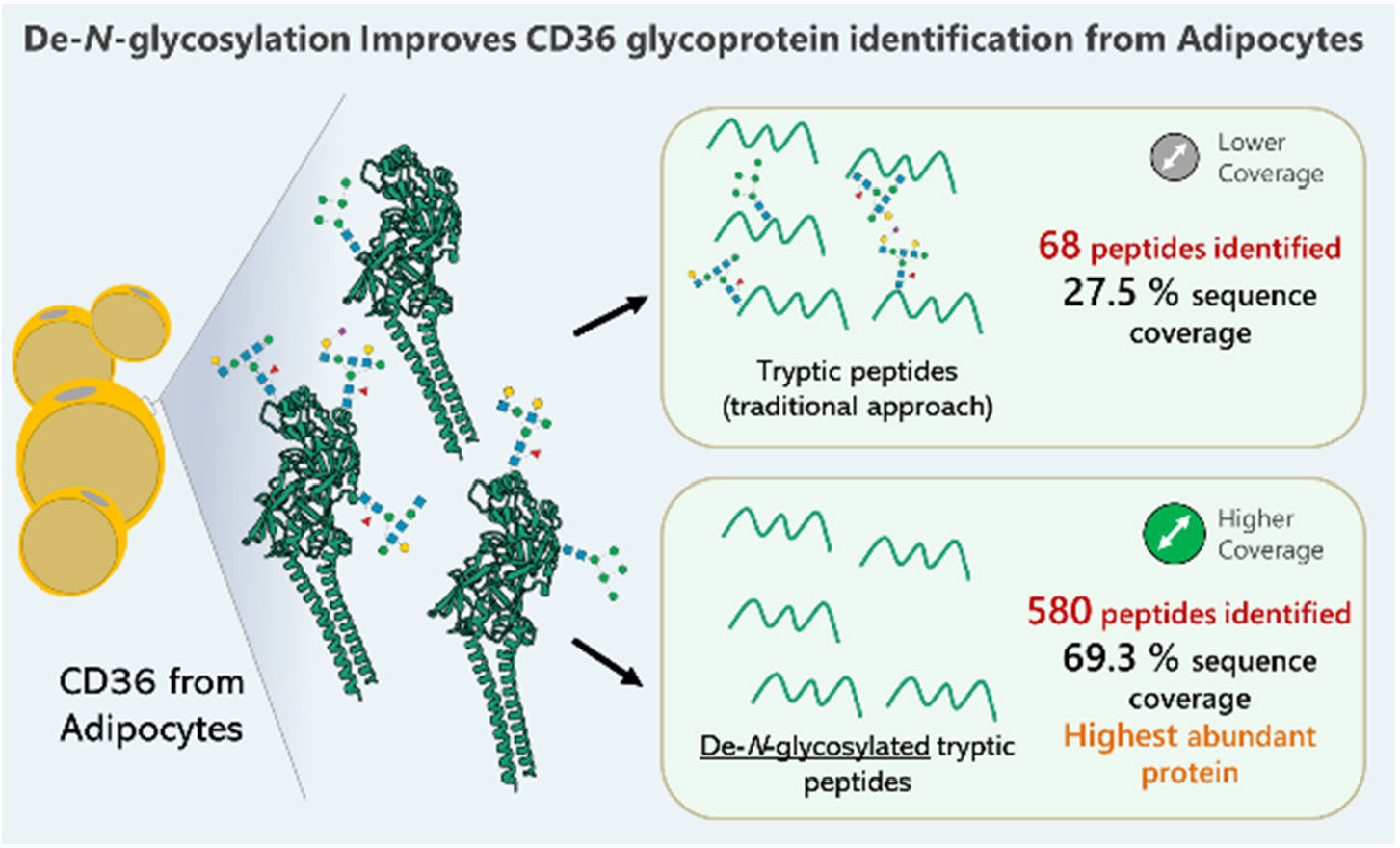

## INTRODUCTION

The cell surface membrane contains a myriad of proteins that initiate a variety of intracellular signalling cascades (1). This protein profile is constantly changing and adapting as part of response and interaction with the external environment. Stem cell differentiation is one such process that influences protein changes on the cell surface as the cell responds to signals from within the stem cell niche. By characterising and quantitating these protein changes during stem cell differentiation, the processes, pathways and networks underlying differentiation may be further understood.

Some studies have investigated the whole proteome during stem cell differentiation (1, 2), but few have specifically targeted the plasma membrane (PM) proteome (3, 4). Much of this effort has been with the aim of identifying new stem cell markers, as one of the current criterion used to identify stem cells is a panel of membrane proteins called ‘clusters of differentiation’ or ‘CD markers’ (5).

Adipogenesis is the process of adipose-derived stem cell (ADSC) proliferation followed by differentiation into mature adipocytes (6). When carried out *in vitro*, this process encapsulates many of the changes seen *in vivo*, making *in vitro* models ideal to study the mechanisms underpinning stem cell differentiation and adipocyte biology (7). Proteomic investigations of adipogenic differentiation in human stem cells in particular have included whole-cell protein analysis of adipogenically differentiated bone marrow-derived mesenchymal stem cells (1) and of adipose-derived stem cell (ADSCs) (8-10). Adipogenic differentiation of ADSCs has also been investigated at the transcriptome level (9, 11) and by membrane CD marker profiling using flow cytometry (12). However, to our knowledge, such an analysis has not been attempted by a targeted membrane proteomics approach where the surface glycoproteins that interact with the stem cell niche can be more readily analysed.

Mass spectrometry (MS) is largely the method of choice for identifying and characterising proteins (13), and when paired with various techniques such as sample-specific preparation, pre-MS sample fractionation methodologies and post-MS bioinformatics tools (14), MS offers a powerful tool to probe the depths of the proteome. Following from an analysis where we previously reported (15) membrane protein *N*- and *O*-glycosylation changes during adipogenic *in vitro* vs *in vivo* differentiation, particularly that bisecting GlcNAc structures are the most abundant structural type in the cell membrane protein *N*-glycome of native adipocytes and characteristic of human adipogenesis, an analysis of the corresponding membrane proteome was undertaken here. Although the function of bisecting GlcNAc carrying structures in adipocytes and stem cells is currently unknown, effects on membrane protein function upon their addition to the *N*-glycan core have been previously described (16-18). The aim here was to investigate other cell surface molecular changes occurring during adipogenic differentiation and, as an orthogonal analysis of protein glycosylation, to characterise the population of proteins from which the glycans analysed previously were released. Cell membrane enrichment using ultracentrifugation and phase-partitioning of hydrophobic proteins was followed by sodium dodecyl sulphate polyacrylamide gel electrophoresis (SDS-PAGE) separation of membrane proteins. By enriching for hydrophobic membrane proteins, this enriched a population of proteins which also included plasma membrane (PM) proteins, allowing changes at the cell surface to be targeted in addition to the global membrane protein analysis (aided by Gene Ontology analysis). Using a label-free quantitative shotgun proteomics workflow, samples were subjected to nanoLC/MS-MS to characterise and compare membrane protein changes during *in vitro* and *in vivo* adipogenic differentiation. Given cell membrane CD markers change upon induction of stem cell differentiation (12), together with the changes in glycosylation we previously observed (15), broader membrane protein changes were explored in cells which had undergone *in vivo* and *in vitro* adipogenesis, by analysis of peptides using reversed phase-liquid chromatography (RP-LC) coupled to ESI-MS/MS. Proteomic comparisons were also performed with and without prior in-gel de-*N*-glycosylation using PNGase F.

## MATERIALS AND METHODS

### Materials

Lithium dodecyl sulphate (LDS) sample buffer, 3-(N-morpholino)propansulfonic acid (MOPS) buffer, NuPAGE Novex 4-12 % (w/v) polyacrylamide Bis-Tris precast gels were purchased from Invitrogen (Life Technologies, Australia). Bradford assay, dithiothreitol (DTT), ammonium bicarbonate (NH_4_HCO_3_), formic acid, acetonitrile (ACN) and iodoacetamide (IAA) were from Sigma (St Louis, MO, USA). μC_18_ ZipTips were from Eppendorf (Hamburg, Germany), and 10 % (w/v) polyacrylamide TGX gel and Coomassie Brilliant Blue G-250 dye were from Biorad (NSW, Australia). HPLC-grade water was obtained using a Milli-Q synthesis water purification system (Millipore, MA, USA). Magic C18 resin was from Michrom Bioresources (Auburn, CA) and sequencing grade modified Trypsin from Promega (WI, USA). Anti-CD36 and anti-rabbit antibodies were obtained from RabMAb (Abcam, Australia).

### Sample collection and membrane protein enrichment

The human abdominal lipo-aspirates used to derive adipose-derived cells were obtained from six patients undergoing elective routine liposuction procedures, and was approved by the Macquarie University Human Research Ethics Committee (Ref: 5201100385) and processed as per our previously reported work (15). Briefly, adipocytes from collagenase-digested adipose tissue were collected in 5 ml amounts for analysis, and the adipose-derived stromal vascular fraction introduced to cell culture to isolate plastic adherent ADSCs. At passage three, half of the culture flasks were harvested for proteomic analysis of the ADSCs while the remaining cells were used for induction of adipogenic differentiation. Cell membrane proteins were isolated from the native adipocytes, ADSCs and dADSCs using Triton X-114 phase partitioning to enrich for hydrophobic membrane proteins as described in (15) as based on Nakano et al. (19).

### SDS-PAGE fractionation of membrane proteins and in-gel digestion

For global shotgun proteomics, membrane proteins from native adipocytes, ADSCs and dADSCs from three different individuals (Individuals 1-3) were quantified using Bradford assay (20). Approximately 25 μg of protein from each sample was reduced with 1 μl 100 mM DTT at 70 °C for 10 min in LDS sample buffer diluted with water for a total volume of 14 μl. IAA (1 μl 500 mM) was added to each sample and proteins alkylated at room temperature for 30 min. Samples were then loaded onto a NuPAGE Novex 4-12% (w/v) polyacrylamide Bis-Tris precast gel and electrophoresed for 2 h at 100 V using MOPS buffer. The gel was fixed with 10 % (v/v) methanol, 1 % (v/v) acetic acid before overnight staining with Coomassie G-250. Stained proteins were visualised using an Odyssey Infrared Imaging System (LiCor).

Each gel lane was excised from the gel and cut into six equal fractions, and then each chopped into smaller pieces which were transferred into Eppendorf tubes for each fraction. Gel pieces were washed twice with 50 % (v/v) ACN and then dehydrated with 100 % (v/v) ACN. This was followed by rehydration with 100 mM NH_4_HCO_3_ for five min before dehydration again with 100 % (v/v) ACN. This was repeated until all gel pieces were de-stained. For protein digestion, 1 μl trypsin (50 mg/μl in 1 mM HCl) was diluted in 2 μl 25 mM NH_4_HCO_3_ and applied to dehydrated gel pieces for 10 min at 37 °C. Approximately 50 μl 100 mM NH_4_HCO_3_ was added to each sample to ensure gel pieces were covered for overnight incubation at 37 °C. Peptides were extracted with 30 μl 25 mM NH_4_HCO_3_ followed by 15 min sonication in a water bath and collected into a fresh Eppendorf tube. Gel pieces were dehydrated with 100 % (v/v) ACN followed by 15 min sonication and the supernatant pooled with the initial extracts. Gel pieces were then washed twice with 5 % (v/v) formic acid followed by sonication and the supernatants collected and pooled at each step. Peptide extracts were then dried using a vacuum centrifuge and resuspended in 10 μl 0.1 % (v/v) formic acid. For desalting of samples, peptides were applied to a μC_18_ ZipTip and the flow through reloaded to maximise recovery. Peptides were eluted in 90 % (v/v) ACN, 0.1 % (v/v) formic acid and dried under vacuum.

### Nanoflow RPLC-MS/MS

Each sample was resuspended in 10 μl 2 % (v/v) formic acid and the peptides separated and analysed using nanoflow LC-MS/MS with a LTQ-Velos Pro linear ion trap mass spectrometer (Thermo, San Jose, CA). Peptides were separated on a reversed-phase column using an Easy nLC II system (Thermo, San Jose, CA). The reversed-phase column was made in-house by packing a fused silica capillary with an integrated electrospray tip with 100 Å, 5 μm Magic C18 resin to approximately 10 cm (with 100 μm inner diameter). A vented pre-column of 3 cm x 100 μm inner diameter packed with the same material was used upstream of the analytical column for sample loading. An electrospray voltage of 1.8 kV was applied via a liquid junction upstream of the C18 column. Samples were injected onto the column followed by an initial wash step with buffer A (5 % (v/v) ACN, 0.1 % (v/v) formic acid) for two minutes at 550 nl/min. At a flow rate of 500 nl/min, peptides were eluted with 0–50 % buffer B (95 % (v/v) ACN, 0.1 % (v/v) formic acid) for 38 min at 500 nl/min followed by 9 min at 800 nl/min. The column eluant was directed into a nanospray ionisation source of the mass spectrometer. Spectra in positive ion mode were scanned over the range *m/z* 400–1500 and, using Xcalibur software (Version 2.06, Thermo), automated peak recognition, dynamic exclusion of 90 s and MS/MS of the top nine most intense precursor ions at 35 % normalisation collision energy were performed.

### Database search for peptide and protein identification

The LTQ-Velos Pro.raw output files were converted into mzXML format and searched against the curated human Uniprot database (20, 264 proteins as of 14^th^ March 2014). This was performed using the Global Proteome Machine (GPM) software (version 2.1.1) using the X!Tandem algorithm (21). The six fractions of each gel-separated membrane protein sample were processed sequentially, generating output files for each fraction along with a merged non-redundant output file for the sample. Peptide identification was determined using a 2 Da parent ion tolerance, 0.4 Da fragment ion tolerance, carbamidomethyl was considered as a complete modification, while oxidation of methionine and threonine, and carbamylation of N-termini and lysine were considered as partial modifications. Reverse database searching was also conducted for estimating false discovery rates (FDRs).

### Data processing and quantitation

For the proteomics results from targeted gel-band analysis, the lists of proteins identified in the membrane fraction of ADSCs, dADSCs and adipocytes were exported from GPM and filtered manually. Contaminations such as bovine serum albumin and porcine trypsin and were removed and only proteins identified with a minimum of two or more unique peptides were retained. After ranking the proteins according to the number of unique peptides, a cut-off was applied after the occurrence of the first reversed hit (false discovery identification) which allowed protein FDR calculation. Proteins were selected as *N*-linked glycoproteins if annotated in the UniProt-KB database (http://www.uniprot.org/).

For the traditional shotgun proteomics approach, the lists of proteins identified at low stringency for three biological replicates of each cell sample were exported from GPM and contaminants removed as above. Protein identifications were further processed using the Spectral Counting Reporting and Analysis Program (Scrappy) (22). For each respective set of three biological replicates for adipocytes, ADSCs and dADSCs, the data were combined to produce a single list of very high stringency identified proteins. These were generated using the following criteria:

a. The protein must be present in all three replicates and,
b. Have a minimum total of five or more peptides across all three replicates in at least one sample type

For example, for a given protein, if the ADSCs had ≥ five total peptides and the dADSCs only three, with no missing values in any of the replicates, it was still considered present in the dADSCs and used for pair-wise comparison. For proteins where the ADSCs had ≥ five peptides and dADSCs had none or missing values in a replicate, it was treated as present in the former and absent in the latter.

Peptide FDR was calculated by 2 x (total number of peptides identified for reversed protein hits/total number of peptides identified for all proteins in the list) x 100 (23) and protein FDR calculated by (number of reversed protein hits/total number of proteins in the list) x 100.

For quantitation, normalised spectral abundance factors (NSAF) were calculated using the following formula: NSAF = (SpC/L)/Σ(Spc/L), where SpC refers to spectral count (number of non-redundant peptide identifications for a given protein) and L refers to the length of the protein (24). Half a spectral count (0.5) was added to all counts to compensate for null values before the natural log of the NSAF values were taken for data normalisation and statistical analysis. Data quality was further examined by generating box plots and kernel density plots of log NSAF values to ensure normal distribution.

### Statistical methods

Student’s *t*-tests were carried out for pair-wise comparisons (Adipocytes vs. ADSCs, and dADSCs vs. ADSCs) and reproducibly identified proteins with *P* < 0.05 and fold change ≥ 1.5 were considered differentially expressed. Analysis of variance (ANOVA) was conducted on proteins shared between ADSCs, dADSCs and adipocytes. Proteins with significantly changing expression (*P* < 0.05) were clustered on a heatmap using a correlation-based distance.

### Gene Ontology annotation

Lists of differentially expressed proteins, proteins that were only identified reproducibly in a given sample, and proteins shared between adipocytes and dADSCs were submitted to the Database for Annotation, Visualisation and Integrated Discovery (DAVID) for functional categorisation (25, 26). Proteins were categorised according to Biological Process at a cut-off of GO level 4, Cellular Component at a cut-off of GO level 3, and using the Kyoto Encyclopaedia of Genes and Genomes (KEGG) pathway analysis tool, with a defined minimum of two proteins in a category. To statistically determine over- or under-represented GO terms in each sample (category enrichment), submitted proteins were compared with the background of total human proteins in DAVID (a default feature of DAVID when no background list of proteins has been also submitted). GO categories that satisfied a test *P*-value < 0.05 in addition to a Benjamini-Hochberg value greater than the test *P*-value were then retained as significantly enriched. The default inclusion of a Benjamini-Hochberg test in DAVID corrects for multiple testing error that may increase the number of false positive identifications (27). The number of proteins in each category was then calculated as a percentage of the number of proteins successfully categorised by DAVID. Similar GO categories that satisfied the statistical criteria outlined above (enriched categories) were grouped using DAVID’s Functional Annotation Clustering analysis at medium level stringency (default level) to reduce redundancy of terms. This analysis uses an EASE score, a modified Fisher Exact *P*-value to determine whether the combined terms in a cluster are enriched with respect to the total human proteome (higher EASE scores denote higher degree of enrichment). In instances where a GO term was not recommended on the European Molecular Biology Laboratory-European Bioinformatics Institute (EMBL-EBI) website to be used for direct annotation, the term was manually removed from DAVID output. For information on individual proteins regarding their biological process, cell component or KEGG pathway involvement, DAVID’s Functional Annotation Table tool was used.

### Orthogonal analysis: Anti-CD36 Western blotting

The human lipo-aspirates used to derive cells for membrane protein extraction, gel separation and Western blotting analysis and qRT-PCR work were from three biological replicates, however these were three different individuals to the ones used for proteomic analysis. Proteins were quantified using Bradford assay and approximately 10 μg membrane proteins was reduced using 100 mM DTT in LDS sample buffer at 70 °C for 10 min before loading onto a 10 % (v/v) polyacrylamide TGX gel. Proteins were separated over 60 min at 200 V before semi-dry transfer to a nitrocellulose membrane using the Turbo blot system (Biorad). The membrane was blocked in 3 % (w/v) skim milk in 0.1% (v/v) Tween-20 in phosphate buffered saline (PBST) for one hour at room temperature before washing twice with PBST. The primary antibody (anti-CD36, rabbit anti-human) was applied at 4 °C overnight at a concentration of 1:1000 in 1 % (w/v) skim milk in PBST. The membrane was subsequently washed three times with PBST before incubation with the secondary antibody (anti-rabbit) at a concentration of 1:10 000 in 3% (w/v) milk in PBST for one hour at room temperature. Finally, the membrane was washed three times with PBST before imaging using the Odyssey Infrared Imaging System (LiCor).

### Orthogonal analysis: CD36 mRNA transcript expression by quantitative real-time reverse transcription polymerase chain reaction (qRT-PCR)

For measurement of CD36 mRNA transcript expression, RNA extraction and qRT-PCR was performed as described in our previous work (15), and was carried out in biological triplicate. Crude RNA was extracted from the three cell types using Trizol reagent and RNA purification was done using the RNeasy Mini Kit. The primer pair used for CD36 were GCAACAAACCACACACTGGG (forward) AGTCCTACACTGCAGTCCTCA (reverse), and for house-keeping gene TATA-box were GCACCACTCCACTGTATCCC (forward) and GCTGCGGTACAATCCCAGAA (reverse).

## RESULTS

Using a label-free shotgun proteomics approach, the membrane proteome of ADSCs was characterized and compared with the corresponding *in vivo* (adipocytes) and *in vitro* adipogenically differentiated products (dADSCs). This work was carried out in biological triplicate, using liposuction samples from the same three individuals previously used for membrane *N*- and *O*-glycosylation analysis and measurement of transcriptomic markers of adipogenesis (15). SDS-PAGE separated membrane proteins were cut into fractions, before trypsin digestion, clean-up and RPLC-MS/MS analysis.

### Data Quality

Data quality was first confirmed by plotting log-transformed NSAF values of the identified proteins as box and kernel density plots (22). The alignment of box plot medians (Supporting Information Figure S1) and histograms of replicates from the same sample tracking close together (Figure 4.2b) indicated that data distribution was approximately normal. The greatest variation was seen in the histograms of dADSC replicates which was also observed at the *N*- and *O*-glycan analysis of samples from the same individuals.

In total, 413 proteins were identified, at an overall protein FDR of 1.21 %. Reversed hits were removed from the list to produce a final list of 408 proteins across the three types of adipose-derived cell samples. Specifically, 175 proteins were identified reproducibly in ADSCs, 237 proteins in dADSCs and 264 proteins in adipocytes. A summary of the number of proteins and peptides identified and false discovery rates is shown in Supporting Info Table S1. The full list of proteins identified and their NSAF and SpC information can be found in Supporting Info Table S2 and S3 respectively). Of the 408 proteins identified, 271 (67.1 %) were categorised as general membrane proteins (all organelle membrane proteins, including the plasma membrane) by DAVID at GO level 3. More specifically, 36.4 % of total proteins were categorised as plasma membrane proteins, thus confirming the Triton X-114 membrane protein enrichment.

### Statistical comparison of shared proteins

An analysis of the distribution of reproducibly identified proteins between the three samples is summarised in Figure 1. One hundred and seventeen proteins were detected only in adipocytes, 72 proteins were only in dADSCs and 34 only in ADSCs. The distribution of proteins between ADSCs and dADSCs is in contrast to their previously analysed *N*-glycan profiles (15) where there were no qualitative differences between the *N*-glycomes of ADSCs and dADSCs. Next, to identify differentially expressed proteins, the 82 proteins detected in all three cell types were compared. Twenty-five proteins were found to be significantly changed (*P* < 0.05) and are visualised on a heatmap with clustering using an ANOVA correlation-based distance (Figure 2). The majority (72%) of these 25 common proteins were identified as membrane proteins, with 60 % categorised as PM proteins (shown in blue font in Figure 2) indicating significant changes occur on the cell surface during adipogenesis. Meanwhile eight of the 25 proteins are glycoproteins as reported in UniProtKB.

**Figure 1.**
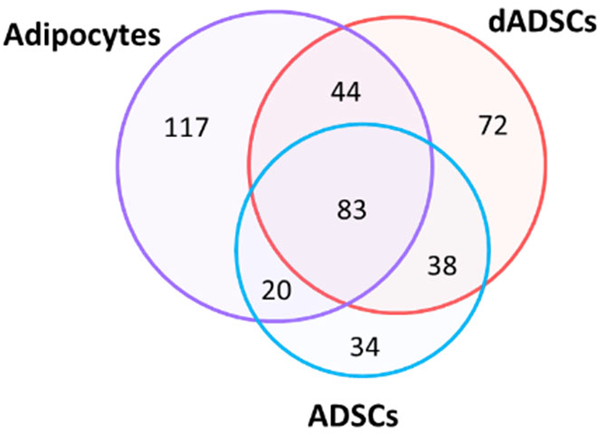
Venn diagram distribution of total proteins identified in the three cell types

**Figure 2.**
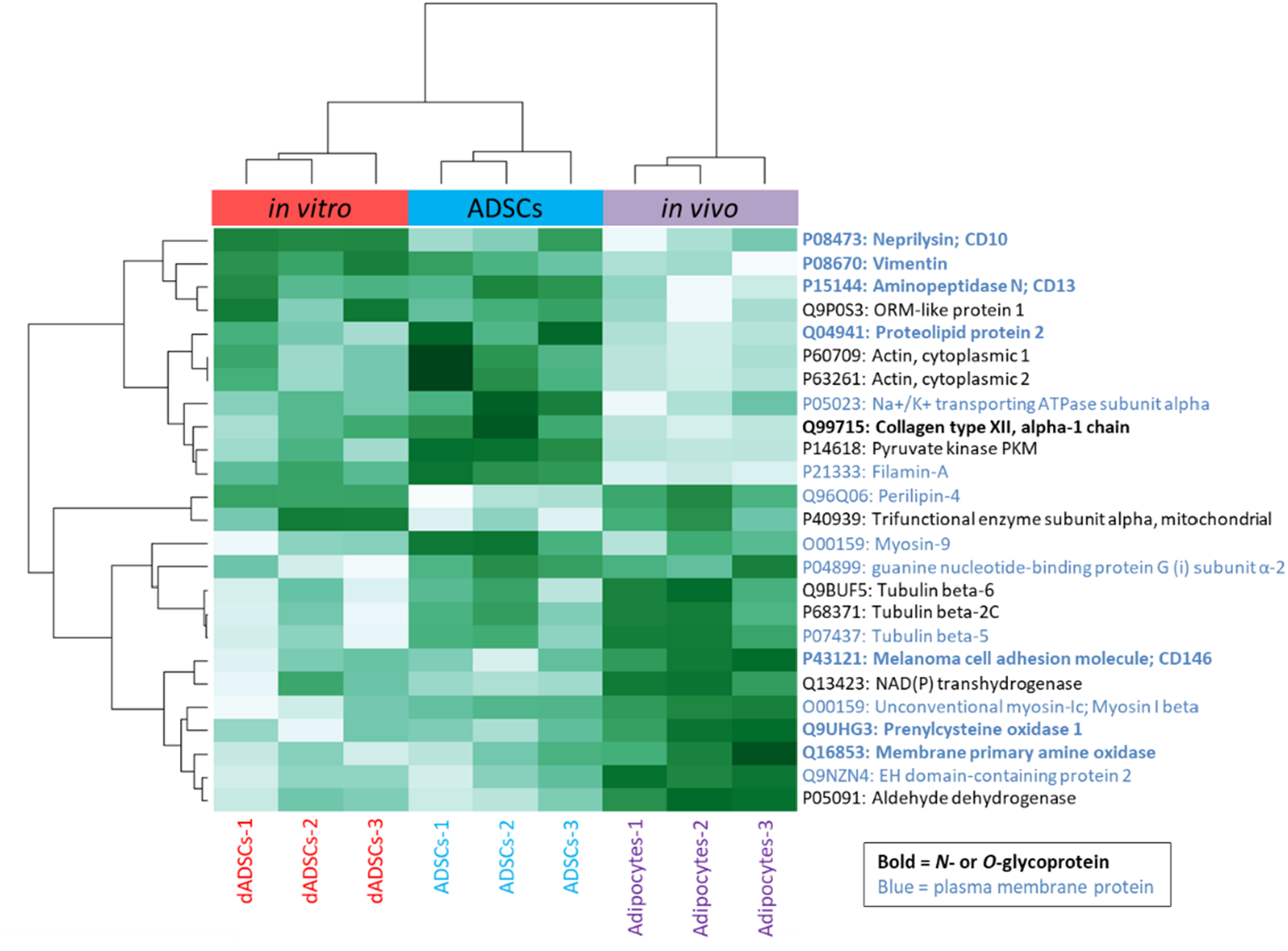
Proteins shared between ADSCs, dADSCs and adipocytes exhibiting differential expression (*P* < 0.05). Proteins are clustered by logNSAF values using a correlation-based distance, and those identified as PM proteins by DAVID are highlighted with blue text while *N*- or *O*-glycoproteins are indicated in bold formatting. Green and white tones indicate up- and down-regulation respectively. The sample name suffixes “-1”, “-2” and “-3” denote the biological replicate number (n=3).

Two proteins in particular displayed higher abundance in both adipogenic products compared to the progenitor ADSCs indicating that their expression during *in vitro* adipogenesis of ADSCs mirrored that of *in vivo* adipogenesis into adipocytes. These proteins were the adipogenic marker, Perilipin-4 (Uniprot: Q96Q06), and a mitochondrial protein, trifunctional enzyme subunit alpha (Uniprot: P40939) (Figure 2). A third protein, myosin-9 (Uniprot: O00159), displayed a lower abundance in both dADSCs and adipocytes when compared to ADSCs. Myosin-9 is classed as an unconventional myosin and has been demonstrated to be involved in insulin sensitive glucose uptake (28). As Perilipin-4 is used as one of the proteins to assess that *in vitro* differentiation has occurred, it is not surprising to see its expression up-regulated. Although it is used as part of a panel of markers, these proteomic results raise the question of whether it is representative enough to be used as a measure of adipogenesis given the number of other dADSCs proteins that did not reflect the levels of their counterparts in the real adipocytes.

A number of cytoskeletal proteins were also identified, namely different types of actins that were down-regulated in dADSCs and adipocytes, while tubulins were up-regulated in adipocytes, highlighting their role in the dynamic changes in cell shape occurring during differentiation. Otherwise, the majority of proteins showed more similarity in abundance between dADSCs and ADSCs than with adipocytes, indicating that the protein abundance changing in *in vitro* adipogenesis does not closely reflect the proteome of the in *vivo* generated mature adipocyte, as shown by broad inspection of the heatmap (Figure 2).

### Pair-wise comparisons of ADSCs with their adipogenic product cells

For a broad overview of the changes in protein abundance occurring between ADSCs and their *in vitro* and *in vivo* adipogenic products, correlation plots of logNSAF ratios for each model of adipogenesis were separately generated (Figure 3). Proteins with differential expression (Student’s *t*-test *P* < 0.05 and a fold change greater than 1.5) are marked in dark blue circles and up- and down-regulated proteins are shown above and below the diagonal respectively. The number of proteins differentially expressed upon native adipocyte membranes is higher than that observed in dADSCs. In addition to this, large deviations in all protein ratios are seen in native adipocytes with many proteins further from the diagonal (Figure 3) than in dADSCs, indicating that ADSCs undergo a greater number and degree of change in protein abundance during *in vivo* adipogenesis than the *in vitro* induced differentiation.

**Figure 3.**
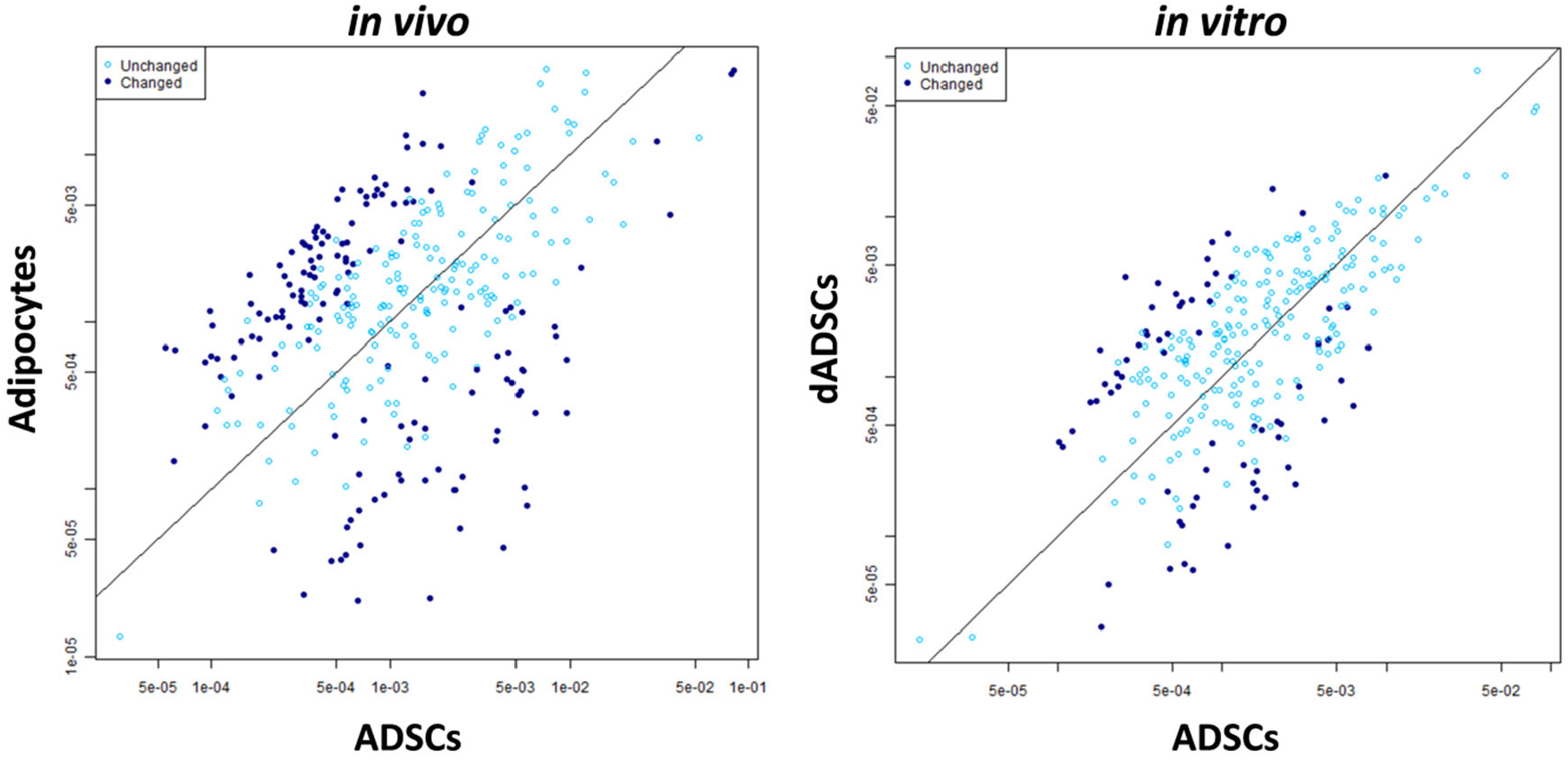
Correlation plots of logNSAF ratios from pair-wise comparison of ADSCs and adipocytes, and ADSCS and dADSCs. Differentially expressed proteins (*P* < 0.05) are marked with a dark blue circle while unchanged proteins are marked with light blue.

Of the five up-regulated proteins in the dADSCs, and nine up-regulated proteins in the native adipocytes, Perilipin-4 and the mitochondrial protein trifunctional enzyme subunit alpha seen in the heatmap were present in both lists (Table 1). Perilipin-4, a structural protein that coats adipocyte lipid droplets (29), was in fact up-regulated to a similar degree in both models with 6.7-fold up-regulation in dADSCs, and 7.0-fold in adipocytes, confirming the abundance pattern observed in the heatmap in Figure 2 and linking this protein directly to adipogenesis. Again, mitochondrial proteins were up-regulated in both *in vitro* and *in vivo* adipocytes and each of these proteins have been previously reported to have roles in both mitochondrial and lipid related processes (30, 31).

**Table 1.**
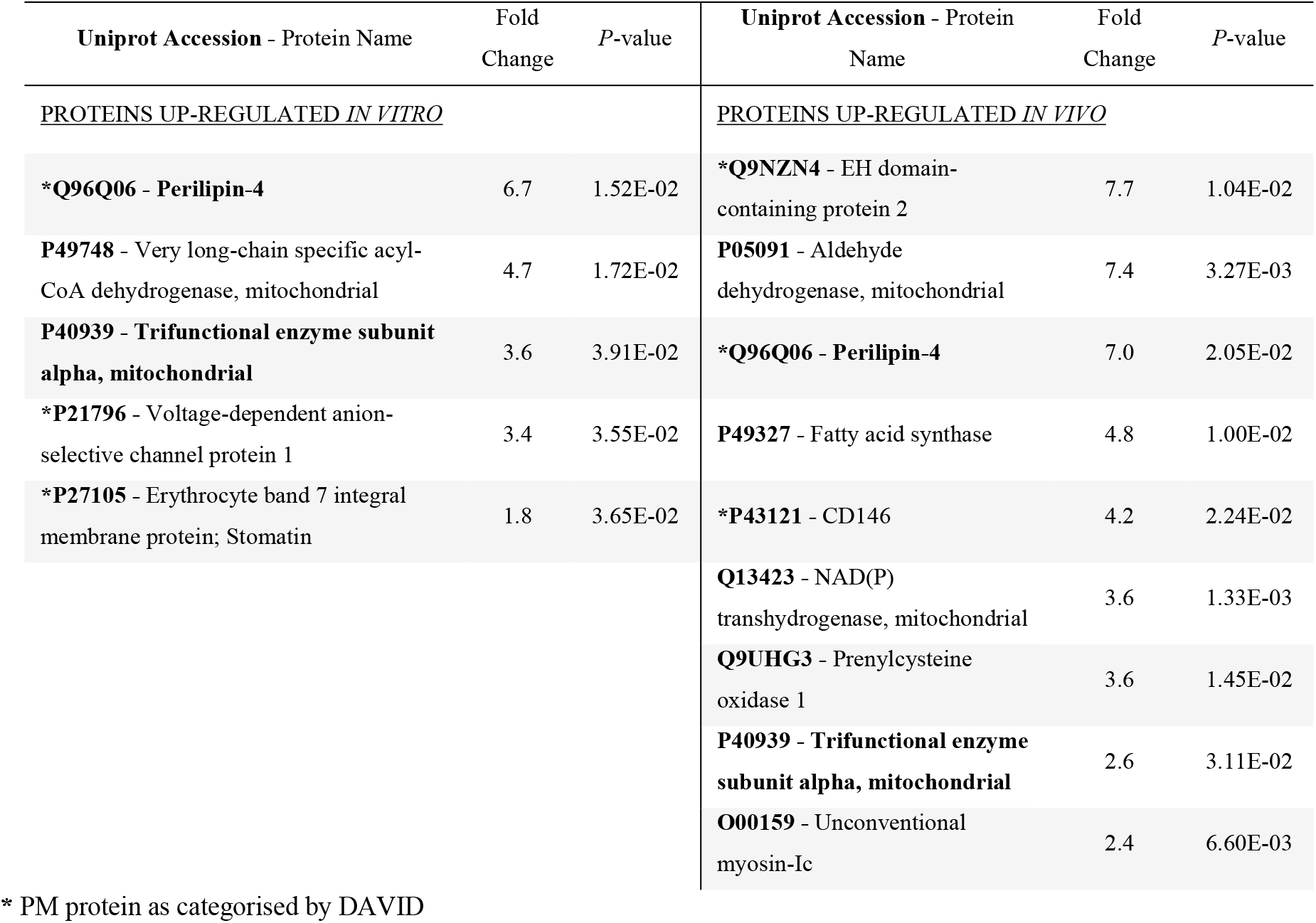
Proteins significantly up-regulated during *in vitro* and *in vivo* adipogenic differentiation based on greatest fold change. The two proteins up-regulated in both adipogenic models are marked in bold.

### Identifying changing glycoproteins in adipogenesis by SDS-PAGE gel

Following the global shotgun proteomic analysis, a targeted approach was taken to try correlate *N*-glycan profiles of different SDS-PAGE separated protein bands with their glycoprotein carriers. In our previous work, an SDS-PAGE separation of adipocyte, ADSC and dADSC membrane proteins from Individual 1 was treated with PNGase F for *N*-glycan profiling and we found the expression of bisecting GlcNAc-carrying glycoproteins were present across almost all the molecular weights of adipocyte SDS-PAGE-separated proteins (15). The follow-up aim here was to attribute this specific glycan structural feature to differentially expressed glycoproteins within a defined molecular weight range (Bands 1-8 on Figure 4).

**Figure 4.**
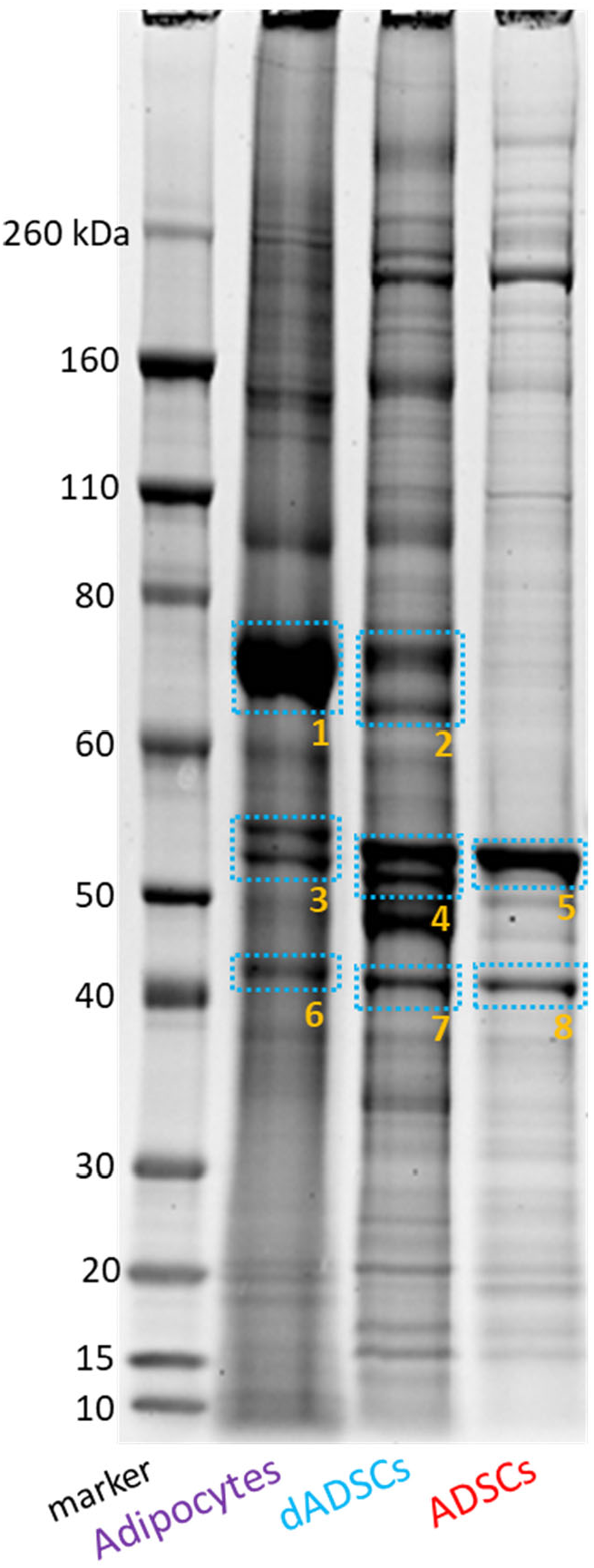
Total membrane proteins separated using SDS-PAGE from native adipocytes, ADSCs and differentiated ADSCs from a representative individual (Individual 1; n=1). Excised gel bands were treated with PNGase F in our previous analysis (15). The proteins identified in each band are listed in Supporting Information Tables S4-11.

Visual differences in protein abundance were particularly obvious between the molecular masses of 60-80 kDa (Bands 1-2) and 50-60 kDa (Bands 3-5), and although equal loading of protein amount was indicated by the Bradford assay, there appeared to be less total protein in the stem cell sample possibly due to the difficulties in solubilizing membrane proteins in protein assay compatible solvents such as the 8 M urea used here. Despite this, the major protein bands are clearly different in the different cell type membrane proteins. Previously PNGase F-treated (de-*N*-glycosylated) protein bands in these mass regions were treated with trypsin before clean up and analysis using RP-LC ESI-MS/MS.

The most striking differences in the protein profiles of the membrane proteins of the three cell types on the gel were at approximately 70 kDa. At this mass region, there was a major protein band (Band 1; Figure 4) in the gel lane containing adipocyte membrane proteins, with two smaller bands present in the dADSC protein lane (Band 2; Figure 4), and little of this molecular weight at all amongst the ADSC proteins. The glycoprotein identified with the highest confidence (lowest log(e) score; Supporting Information Table S4) in Band 1, and in fact the most abundant protein identified, was CD36 glycoprotein, identified with 56 unique peptides and a total of 580 peptides. CD36 is a marker of the adipogenic lineage (32) and was also present in the corresponding region in dADSCs (Band 2; Figure 4) however at much lower abundance compared to native adipocytes with 15 unique and 69 total peptides (Supporting Information Table S5). This was the major protein difference observed at this molecular mass region, translating to a differential expression of CD36 with 9 times the number of total peptides and 3.7 times the number of unique peptides found in the native adipocyte membranes (Band 1) compared with differentiated ADSCs (Band 2). This difference between the two cell types was not apparent using the traditional shotgun total proteomics approach.

Although CD36 has a predicted amino acid sequence molecular weight of approximately 53 kDa, its ten potential *N*-glycosylation sites can result in a predicted weight of up to 90 kDa (33), indicating that the CD36 detected here in adipocytes and dADSCs is highly glycosylated as also indicated by the typical thickness of a heterogeneous glycoprotein gel band. Although CD36 was present in the 70 kDa gel region in dADSCs (Band 2; Figure 4), the most abundant glycoprotein identified in that band was CD73 (Supporting Information Table S5), which is a mesenchymal stem cell marker. The identification of a stem cell marker here indicates that the dADSCs share characteristics of both mature adipocytes and their progenitor stem cells, as also observed in previous *N*- and *O*-glycomic, and transcriptomic analyses of this *in vitro* adipogenic model (15). Further CD36 variants with molecular weights lower than 53 kDa can also be produced from alternate splicing events, and were also detected in the adipocyte gel lane in both the lower 50-60 kDa and ∼40 kDa molecular mass regions (Supporting Information Table S6 & S9).

As seen in Table 2, other improved protein identifications were made if proteins were de-*N*-glycosylated before trypsin digestion. Galectin-3-binding protein (in Band 2) was not detected when the sample was not de-*N*-glycosylated whereas 6 unique peptides were seen after de-glycosylation. In fact, the majority of glycoproteins identified in the de-*N*-glycosylated Band 2 sample were not detected at all in the corresponding glycosylated sample. Glycoproteins identified in both dADSCs and adipocyte cell membranes (Bands 3 & 4, respectively; Fig 4 and Supporting Information Tables S6-7) were prenylcysteine oxidase 1, serpin H1 and calreticulin, whilst adipocyte plasma membrane associated protein was found in all three cell types. Glycoprotein identifications were again improved with the de-*N*-glycosylation of samples, with proteins such as Haemoglobin subunit beta (Supporting Information Table S12) and protein-lysine 6-oxidase (Supporting Information Table S13) not detected in the samples that were not de-*N*-glycosylated.

**Table 2.**
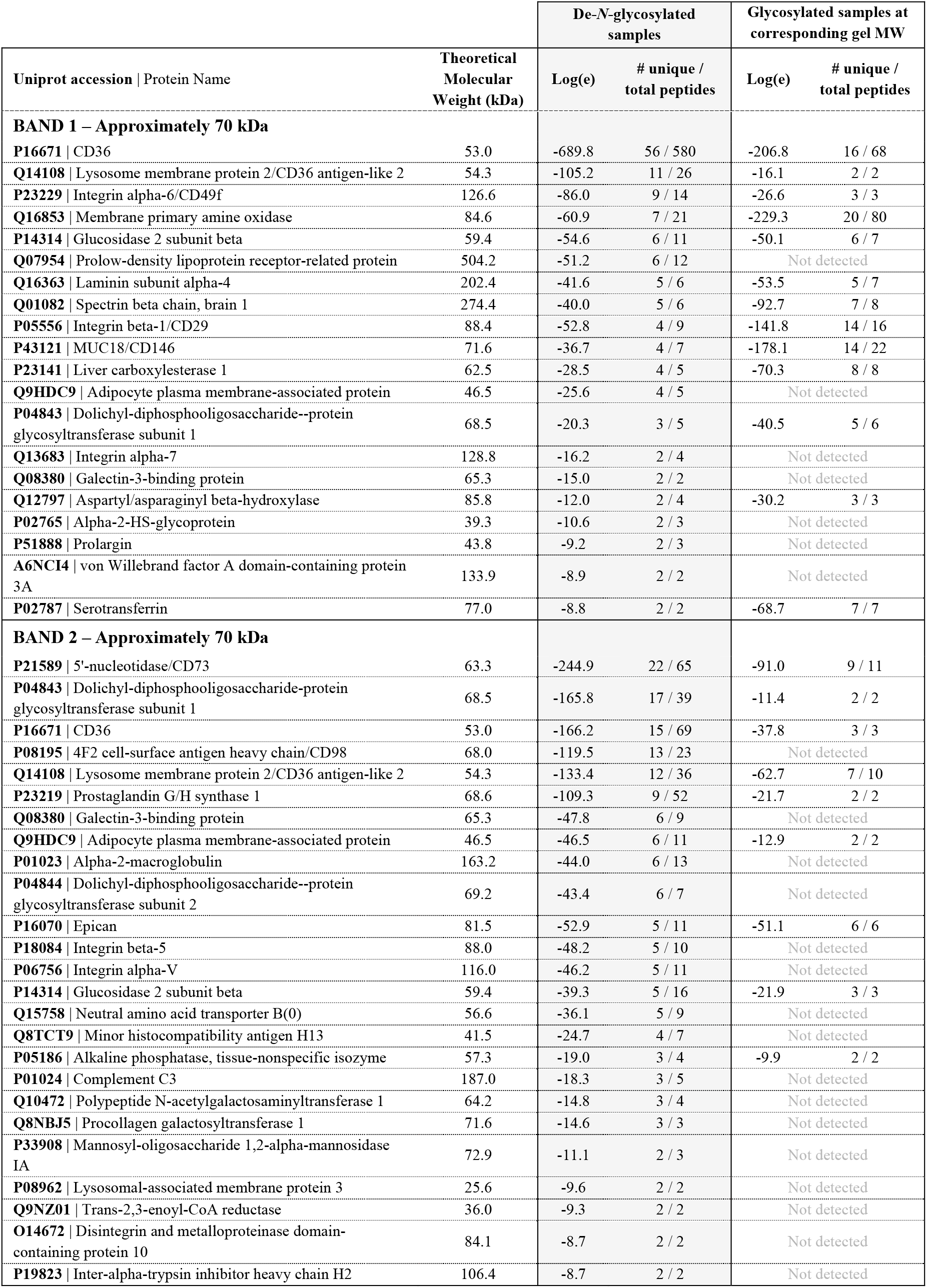
All glycoproteins identified from the targeted proteomics of de-*N*-glycosylated SDS-PAGE adipocyte protein Bands 1 and 2 (Figure 4) listed in order of highest confidence (log(e) score), compared with the corresponding (glycosylated) gel region of the same individual analysed using traditional shotgun proteomics.

The 40 kDa mass region showed a major band in all three cell types, with relatively high abundance in the ADSCs (Figure 4). CD36 was again identified in the adipocytes at this mass range (Band 6, Figure 4) with much better coverage of the protein after de-*N*-glycosylation, but is most likely a cleavage product as the non-glycosylated mass of CD36 is 53 kDa. Again, several glycoproteins in Band 7 specifically were detected in the de-*N*-glycosylated samples and not detected in the glycosylated samples, namely HLA class I histocompatibility antigen B-53 alpha chain, synaptophysin-like protein 1, inositol monophosphatase 3, cation-dependent mannose-6-phosphate receptor, pentraxin-related protein PTX3 and inhibitor of nuclear factor kappa-B kinase-interacting protein (Supporting Information Table S14).

### De-N-glycosylation is essential for detecting accurate CD36 expression

Of all the membrane glycoproteins identified in the targeted analysis, CD36 displayed the most striking differential expression across the three cell types, both visually on the gel (Figure 4) and in the number of unique and total peptides identified between adipocytes and dADSCs (Table 1) – most evident only after de-*N*-glycosylation of the sample. Given CD36 is the top protein identified also in the major protein band of SDS-PAGE separated adipocyte proteins (Band 1, Figure 4), its absence as a differentially expressed protein in the heatmap ANOVA analysis (Figure 2) and amongst the up-regulated proteins in adipocytes (Table 1) in the global shotgun proteomics data was probably due to the inaccessibility of the protein to complete digestion without de-*N*-glycosylation.

As the primary difference between the targeted analysis of the SDS-PAGE (de-*N*-glycosylated) protein bands and the global shotgun proteomics analysis had been the removal or presence of *N*-glycans before trypsin digestion, the CD36 sequence coverage by MS for the de-*N*-glycosylated CD36 and glycosylated CD36 samples were compared (Figure 5 & Table 2). This revealed that only 68 total peptides were identified in the glycosylated sample using traditional shotgun proteomics, whilst 580 were identified when CD36 was de-*N*-glycosylated prior to trypsin digestion, resulting in a 41.74 % increase in sequence coverage (shown in red in Figure 5). The majority of the protein sequence in the glycosylated sample was undetected (highlighted in green). CD36 contains ten *N*-glycosylation sites (N-X-S/T; where X can equal any amino acid except proline), of which nine are found in the undetected green regions (Figure 5A) which strongly indicates that the presence of *N*-glycosylation on the protein hindered the ability of trypsin to cleave the protein and/or prevented the matching of detected glycopeptide MS/MS spectra with theoretical *in silico* peptide sequence masses, thereby lowering the number of identified peptides matching the protein in question.

**Figure 5.**
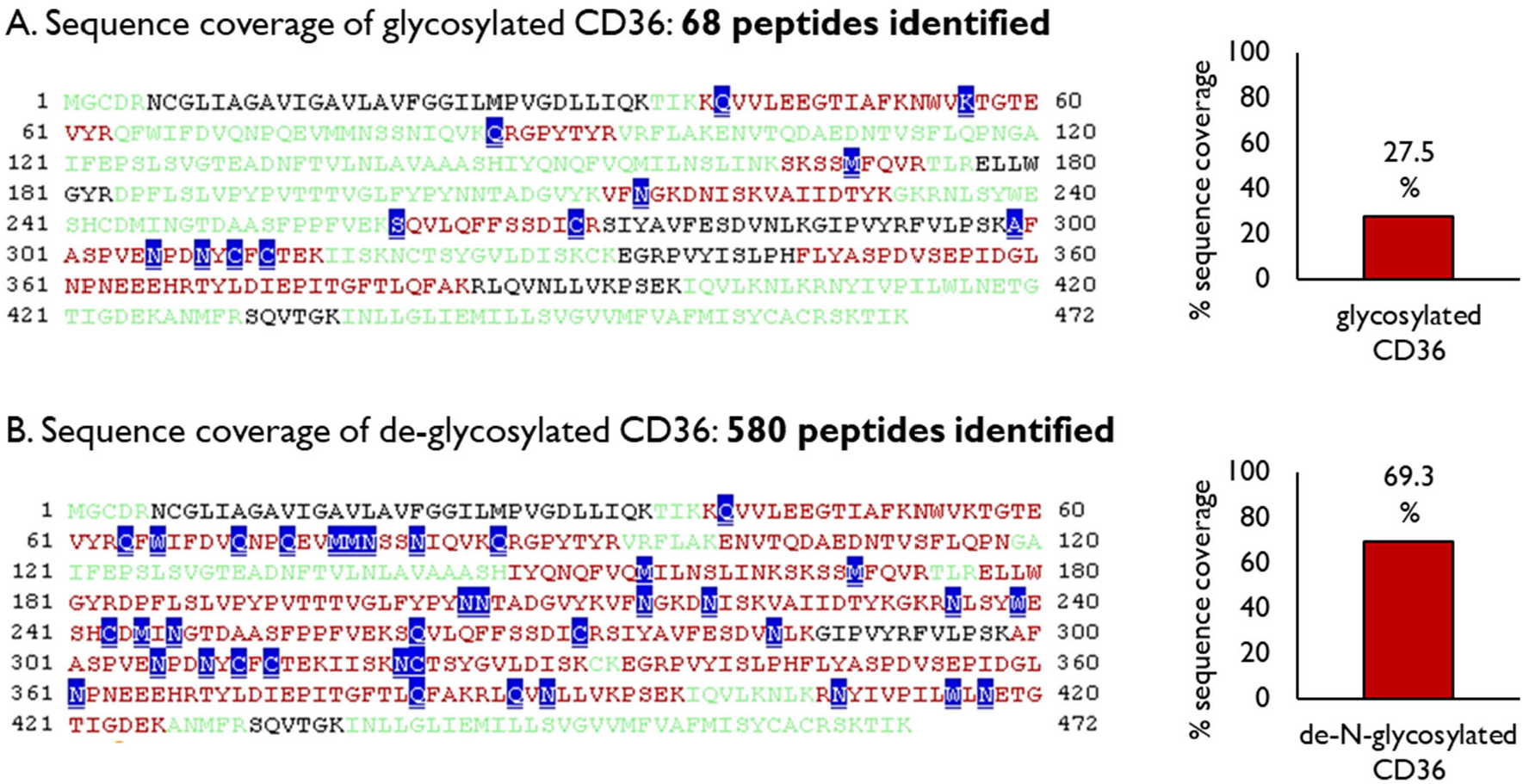
CD36 protein sequence coverage in the Global Proteome Machine using the X!Tandem algorithm for peptide MS/MS spectra matching with *in silico* fragments with (**A**) *N*-glycans present (27.54 % sequence coverage) and (**B**) removed (69.28 % sequence coverage) prior to trypsin digestion. Green amino acid regions indicate difficult to match or unmatched areas, while red regions indicate areas matched with the *in silico* proteome. Chemically modified residues are shown in blue.

To confirm the differential expression of CD36 during *in vivo* and *in vitro* adipogenesis, and validate it as the most abundant protein in adipocytes as unearthed through proteomics of de-*N*-glycosylated protein gel Bands 1 and 2, CD36 expression was further measured using two orthogonal approaches. Western blotting with an anti-CD36 antibody of the gel-separated membrane proteins and qRT-PCR measurement of the CD36 gene transcripts was conducted in biological triplicate. The results of the Western blot displayed binding of the anti-CD36 antibody only in the 70 – 80 kDa molecular weight range, with the strongest signal observed in adipocytes followed by dADSCs, and almost absent in ADSCs (Figure 6a) supporting the mass spectrometric abundance analysis of de-*N*-glycosylated CD36 combined with visual inspection of the gel. While no bands were seen in the 40 kDa molecular range, despite identification of CD36 by MS/MS at this mass in adipocytes and dADSCs (Supporting Information Table S14), this may be due to the antibody not recognising this isoform. The region of CD36 recognised by the antibody is between amino acids 350-450 which contains one *N*-glycosylation site motif at Asn^417^ such that the detection of CD36 by MS/MS at the 40 kDa range is most likely a cleavage product containing an area of the protein not recognised by the antibody.

**Figure 6.**
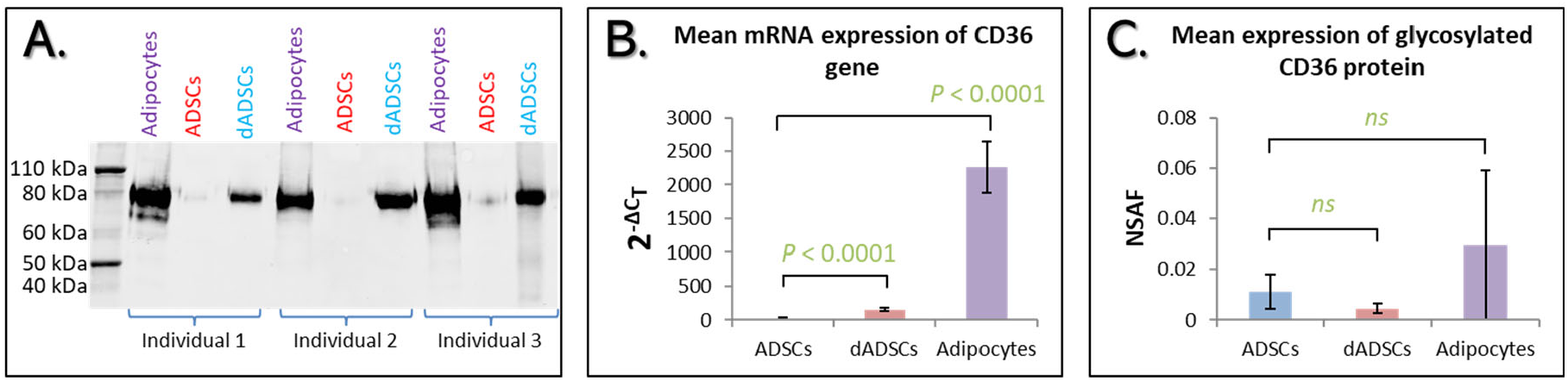
(**A**) Western blot probing for CD36 in adipocyte, ADSC and dADSC membrane proteins in three biological replicates. Comparison of the mRNA transcript (**B**) and shotgun proteomic expression levels (**C**) of the adipogenic marker CD36 in its glycosylated form. For transcript analysis, expression levels were normalised to the TBP housekeeping gene and are shown as 2^−ΔCT^ mean ± standard deviation. *P*-values from student’s *t*-test comparison of ADSCs and dADSCs, and ADSCs and adipocytes are shown. *‘ns’ =* not significant.

The expression pattern of CD36 observed by Western blotting was also seen in analysis at the mRNA transcript level. The qRT-PCR measurement of CD36 transcripts showed significantly higher expression in both mature adipocytes (102.9-fold increase) and dADSCs (7.1-fold increase) compared to ADSC progenitors (Figure 6B). Expression was highest in adipocytes as was observed by Western blot at the protein level (Fig 6A). This finding supports the proteomic analysis of de-*N*-glycosylated CD36 from gel Bands 1 and 2 (Figure 4) and is in contrast to the global shotgun proteomic analysis of these samples (Fig 6C), which did not reveal any significant differences in the abundance of the CD36 protein between ADSCs and both adipogenic products (Figure 1C), despite CD36 being a known adipogenic marker.

Together, the results of both techniques carried out at the different gene-product levels (mRNA and antibody detected protein expression), indicate that CD36 protein abundance was masked in the global shotgun proteomic analysis when de-*N*-glycosylation of the proteins was not carried out prior to MS analysis. This highlights the crucial need for de-*N*-glycosylation prior to proteomics analyses regardless of whether the experiment has a glycomics-based aim, to allow the proper detection and identification of glycoproteins and their abundance. The results here have also demonstrated the value of verifying protein expression patterns using orthogonal techniques.

## DISCUSSION

In this study, total membrane proteins from native adipocytes, ADSCs and *in vitro* differentiated ADSCs (dADSCs) from three liposuction patients were separated using SDS-PAGE and investigated using a label-free quantitative shotgun proteomics approach. A total of 408 proteins were reproducibly identified across the three cell types, of which 67.1% were membrane proteins which is in good accordance with the percentage membrane protein enrichment seen in previous studies

A greater degree of change was observed in the proteome during *in vivo* differentiation compared to *in vitro* differentiation, mirroring the results of the global *N*-glycan profiling from cells from the same individuals (15). This indicates that changes in the stem cell glycosylation machinery towards an adipogenic profile are either slower during 30-day *in vitro* adipogenesis compared to protein expression changes (e.g. induction of CD36 expression in dADSCs). In the pair-wise comparison of ADSCs and their differentiated adipogenic progeny, the majority of proteins were largely unchanged, which is surprising given the drastic morphological changes associated with adipogenic differentiation. The two proteins that were up-regulated in both *in vitro* and *in vivo* adipogenic products were the mitochondrial trifunctional enzyme subunit alpha and Perilipin-4. Trifunctional enzyme subunit alpha has also been reported to be up-regulated in adipogenically differentiated mouse fibroblasts compared to undifferentiated fibroblasts (34) and is involved in catalysing long-chain fatty acid oxidation in the beta-oxidation pathway (31). Perilipin-4 is a described adipogenic marker because of its involvement in lipid vesicle structure in adipocytes (29, 30) therefore its up-regulation in adipocytes is expected, and the significant increase of Perilipin-4 protein in dADSCs in the present study suggests its inclusion as a marker to evaluate *in vitro* differentiation of stem cells into adipocytes is warranted.

Following the observed changes in the global membrane protein *N*-glycosylation of stem cells during *in vitro* and *in vivo* adipogenesis and in the membrane proteome in this study, an integrated gel-based glycomics and proteomics approach was undertaken to further investigate specific differences between the cell types. Membrane proteins from native adipocytes, ADSCs and *in vitro* differentiated ADSCs from a representative individual were previously separated using SDS-PAGE (15) and the separated protein bands were analysed by *N*-glycomic profiling. Here, complementary shotgun proteomics was performed on the leftover de-*N*-glycosylated protein bands to try determine the correlation between expressed proteins and their glycosylation profile. The proteomic analysis of de-*N*-glycosylated membrane proteins separated by SDS-PAGE revealed different sets of glycoproteins compared to the corresponding total PM samples not treated prior with PNGase F. This approach revealed that the adipogenic marker CD36 is the major (glyco)protein in adipocytes and is also expressed in dADSCs upon adipogenic differentiation. This observation was confirmed using orthogonal techniques of Western blotting and mRNA expression of CD36.

De-*N*-glycosylation of proteins prior to proteomic analysis has previously been described to improve identification of *N*-glycosylated proteins in shotgun experiments (35-37). Given that CD36 contains up to ten *N*-glycosylation sites (33), removal of *N*-glycans was particularly crucial and resulted in an increase from 27.54 % to 69.28 % in sequence coverage and a higher number of peptides identified. Nine of these *N*-glycan sites were present in regions of the protein that were not identified in the glycosylated sample analysis, most likely due to either incomplete cleavage around the glycosylation site or to the discrepancy between any detected glycopeptide masses and the *in silico* peptide mass matching. *N*-glycans have been reported to obscure the action of proteases such as trypsin and AspN (3, 35). Lower MS ionisation of glycopeptides compared to un-modified peptides (38) and glycopeptide site heterogeneity resulting in a lower copy number of each peptide (39) may also account for the decreased coverage of the glycosylation CD36.

For CD36, the use of Western blotting and measurement of transcript levels served a dual purpose to confirm expression at the protein level between 60-80 kDa as well as the differences observed between the samples. CD36 abundance was confirmed as being highest in native adipocytes by both orthogonal techniques, was induced by differentiation and was significantly higher than the ADSCs at the transcript level. Overall the data confirm the crucial need for sample de-*N*-glycosylation for accurate identification of glycoprotein abundance by classic shotgun proteomics methods, and the value in orthogonal methods to validate differential protein expression. This observation has also been reported by Bailey and Schulz (35) and has previously been employed sparingly in previous proteomics studies (3, 40, 41).

As CD36 and bisecting GlcNAc structures were both up-regulated in the adipogenic cell products, further glycoproteomics-based investigations should aim to determine whether they share a dual functional role in adipogenesis and the unique biology of adipocytes. There is a high likelihood that CD36 carries bisecting GlcNAc-type *N*-glycans, as the *N*-glycan profile of the 60-80 kDa mass region where CD36 was identified contained several bisecting GlcNAc type glycans (15). While CD36 was also identified in the 60-80 kDa mass region in dADSCs the corresponding *N*-glycan MS profile of this region showed no masses corresponding to potential *N*-glycans, indicating that while CD36 protein expression is increased during *in vitro* adipogenesis, its proteo/glyco-forms are not the same as CD36 in a native adipocyte.

In regard to stem cell differentiation studies in general, the strength of this work was that it allowed the comparison of *in vitro* differentiated adipogenic cells with their *in vivo* counterparts (native adipocytes). This comparison, performed by previous membrane protein glycosylation analysis and complementary shotgun membrane proteomics of de-*N*-glycosylated samples here revealed that the *in vitro* generated cells express features that reflect native adipocytes, but also still express stem cell features such as the expression of stem cell marker CD73, indicating incomplete differentiation and a mixed population of still ‘stem-like’ dADSCs alongside more matured dADSCs in the culture flask.

Overall, the work described here presents important analytical considerations when investigating the (glyco)proteome. Most importantly, the results demonstrate that the de-*N*-glycosylation of samples prior to proteomics analysis is an important and crucial consideration for the accurate measurement of relative expression of all proteins, and that not including this step results in under-representation glycoprotein abundances. Given approximately 73% of the human proteome is potentially *N*-glycosylated, this is an essential consideration for studying the proteome. Specifically for CD36, while it is categorised as an adipocyte marker, it is also found on other cell types and is known to be involved in several other biological processes (e.g. cancer progression (42), platelet function and lipid uptake (43)). Its potentially masked abundance in proteomic datasets, as seen here, will naturally obstruct its analysis in other experimental settings and the critical step of de-*N*-glycosylation prior to proteomics analysis therefore extends beyond adipocyte biology.

## ASSOCIATED CONTENT

Supporting Information Section: Supplementary Figure S1, and Supplementary Tables S1-15

## AUTHOR INFORMATION

### Author Contributions

Experiments were designed by KWK, BRH, PAH and NHP. Experiments were performed by KWK, and manuscript written by KWK and NHP. All authors reviewed and approved the final manuscript.

## ACKNOWLEDGEMENTS

The authors would like to thank Dr Sinead P. Blaber and Dr Hannes Hinneburg in guiding the establishment of ADSC culture lines and differentiation experiments. We would also like to thank Dr Anahit Penesyan and Dr Deepa-Ruth Varkey for technical assistance with glycosyltransferase expression profiling using qRT-PCR, and Dr Ling Yen Lee for guidance with Western Blotting.

This research was facilitated by access to the Australian Proteome Analysis Facility, established under the Australian Government’s National Collaborative Research Infrastructure Scheme. K.W-K was supported by an Australian Postgraduate Award. This research was funded partially by the Australian Government through the Australian Research Council Centre of Excellence in Synthetic Biology (Project ID CE 200100029).

The authors have declared no conflict of interest.

